# Microbiota colonization tunes the antigen threshold of microbiota-specific T cell activation in the gut

**DOI:** 10.1101/2022.07.29.501979

**Authors:** Daniel Hoces, Basak Corak, Anna Estrada Brull, Sara Berent, Erica Faccin, Claudia Moresi, Tim Keys, Nicole Joller, Emma Slack

## Abstract

Harnessing the potential of commensal bacteria for immunomodulatory therapy in the gut requires the identification of conditions that modulate immune activation towards incoming colonizing bacteria. In this study, we used the commensal *Bacteroides thetaiotaomicron (B.theta)* and combined it with *B.theta*-specific transgenic T cells, in the context of defined colonization of gnotobiotic and immunodeficiency mouse models, to probe the factors modulating bacteria-specific T cell activation against newly colonizing bacteria. After colonizing germ-free (GF) and conventionally raised (SPF) mice with *B.theta,* we only observed proliferation of *B.theta*-specific T cells in GF mice. Using simple gnotobiotic communities we could further demonstrate that T-cell activation against newly colonizing gut bacteria is restricted by previous bacteria colonization in GF mice. However, this restriction requires a functional adaptive immune system as *Rag1^-/-^* allowed *B.theta*-specific T cell proliferation even after previous colonization. Interestingly, this phenomenon seems to be dependent on the type of TCR-transgenic model used. *B.theta*-specific transgenic T cells also proliferated after gut colonization with an *E.coli* strain carrying the *B.theta-specific* epitope. However, this was not the case for the SM-1 transgenic T cells as they did not proliferate after similar gut colonization with an *E.coli* strain expressing the cognate epitope. In summary, we found that activation of T cells towards incoming bacteria in the gut is modulated by the influence of colonizing bacteria on the adaptive immune system of the host.

## INTRODUCTION

The gut harbors the highest concentration of bacteria present in our body, reaching densities up to 10^10^ to 10^11^ CFUg^-1^ of content[1]. Although most gut colonizing bacteria are excluded from immune cells by a series of barriers such the mucus layer and epithelium[2], there is clear evidence that a fraction of the microbiota is sampled and presented to T cells, inducing immune activation and proliferation without tissue damage[3]. Commensals interact with the immune system, for example by imprinting a particular phenotype on T cells[4],[5] and regulate the balance between health and disease states[6].

Although there is a clear potential for immunomodulatory therapy based on microbiota engineering, we still lack fundamental knowledge on the immunity-microbiota balance. For example, it is assumed that the microbiota constantly calibrates the immune threshold of activation, which then promotes protective immunity against infectious agents[7]. In addition, certain commensals can induce immune activation of antigen-specific T cells and their differentiation into regulatory T cells (Treg)[8],[9]. However, the commensal antigenic load needed to initiate T cell activation in the gut microenvironment is not known. Understanding the modulation of these antigenic thresholds is important as it would serve as a goal for any microbiota engineering therapy either based on the insertion or deletion of a particular immunomodulatory microbe into the community.

In this study, we combined bacteria-specific TCR-Tg T cells with bacteria mutants, gnotobiotic and immunodeficiency mouse models to probe the antigenic threshold of activation for microbiota-specific T cells in the gut under homeostatic conditions. We found that T cell activation and proliferation in the gut is controlled by the combination of the bacterial chassis carrying the antigen, previous bacteria colonization and a functional adaptive immune system.

## RESULTS

### BΘOM T cells proliferation is restricted in colonized mice

To explore the host conditions under which *B.theta* can induce antigen-specific immune activation during gut colonization, we used T-cell receptor transgenic (TCR-Tg) T cells that recognize an outer membrane protein of *B.theta* (BΘOM T cells)[10]. We colonized germ-free (GF) and conventional (SPF) mice with *B.theta* before transferring CD45.1+ BΘOM T cells labelled with a CellTrace Violet (CTV) (Fig.1A). Seven days later, we identified the activated BΘOM T cells in the spleen and mLN (Fig.1B, Suppl.Fig.1).

**Figure 1.**
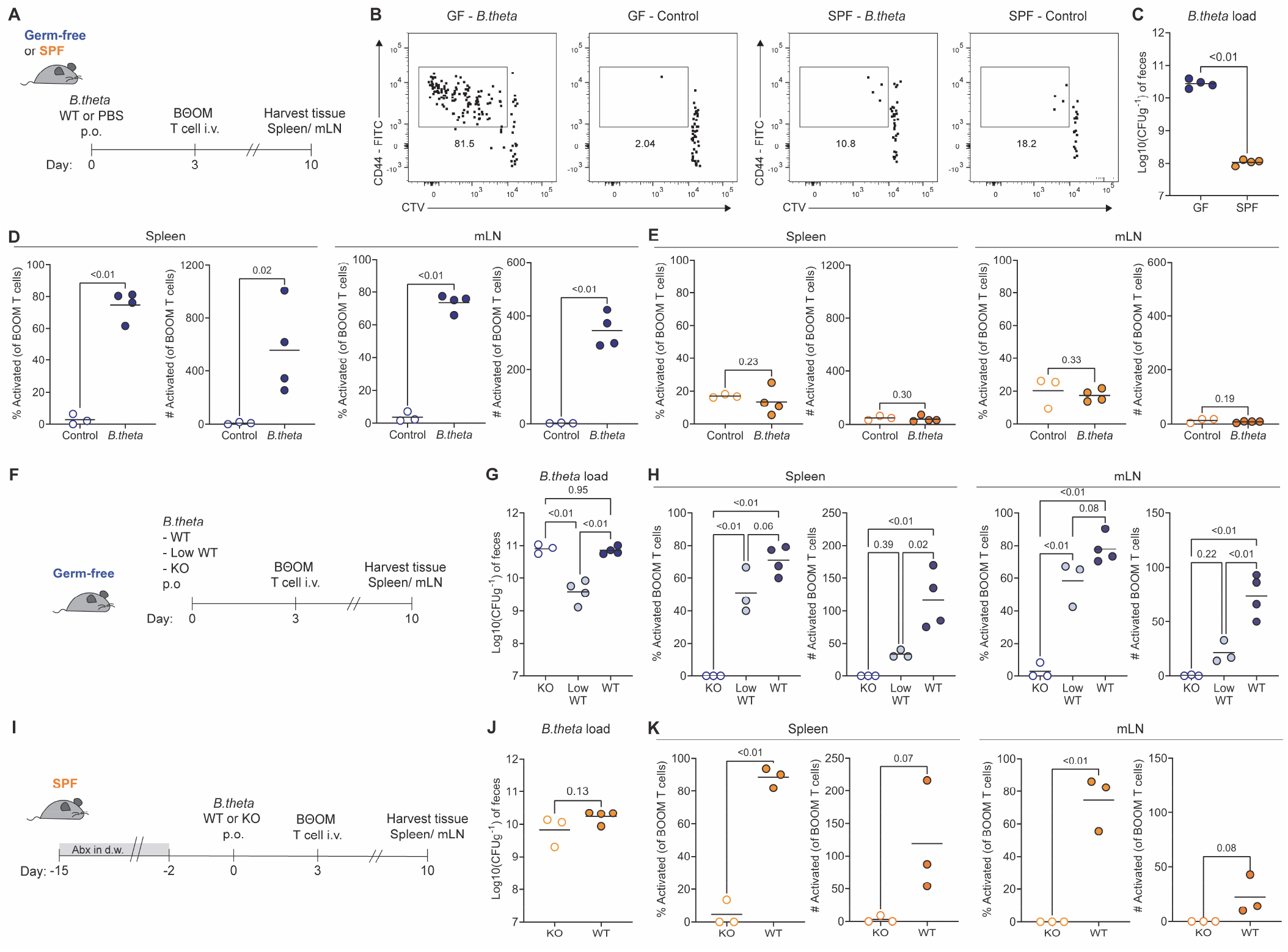
BΘOM T cell proliferation is restricted in colonized mice. **(A)** Experimental set-up of *B.theta* colonization (or PBS control) and adoptive transfer of BΘOM T cells into GF or SPF C57BL/6J mice. **(B)** Representative flow cytometry plots of activated (CD44^hi^ CTV^low^) of BΘOM T cells in GF and SPF mice. **(C)** *B.theta* gut luminal load in GF and SPF mice at the time of BΘOM T adoptive transfer. **(D-E)** Percentage and number of activated BΘOM T cells in spleen and mLN in **(D)** GF and **(E)** SPF mice. **(F)** Experimental set-up of *B.theta* colonization in GF mice with different loads of wild-type *B.theta* strain (WT: all wild-type bacteria; Low WT: one wild-type bacteria per 10 KO bacteria; KO: all KO bacteria). **(G)** Gut luminal load of *B.theta* strains in GF mice at the time of BΘOM T adoptive transfer. **(H)** Percentage and number of activated BΘOM T cells in spleen and mLN. **(I)** Experimental set-up of *B.theta* colonization after broad-spectrum antibiotic treatment. SPF mice were colonized with either wild-type *B.theta* or a mutant strain lacking the BT4295 antigen (KO). **(J)** Gut luminal load of *B.theta* strains in antibiotic-treated SPF mice at the time of BΘOM T adoptive transfer. **(K)** Percentage and number of activated BΘOM T cells in spleen and mLN.

In GF mice, *B.theta* colonization (Fig.1C) significantly increases the percentage of BΘOM T cells that divided, become activated and expand in the spleen and mLN compared to control mice (Fig.1D). However, in SPF mice, there was no significant effect of *B.theta* colonization on BΘOM T cell proliferation (Fig.1E). This differential activation of BΘOM T cells by *B.theta* colonization in GF compared to SPF mice is not observed when mice are challenged systemically (Suppl.Fig.2A), as BΘOM T cells show a similar activation pattern in both GF and SPF mice (Suppl.Fig.2B).

One hypothesis for restriction of T cell proliferation in SPF mice is that the lower level of *B.theta* colonization in the gut (Suppl.Fig.3A) may also reduce bacterial translocation, and therefore antigen availability for BΘOM T cells in gut-draining lymphoid tissues. Although presence of live bacterial colony-forming units (CFU) was highly variable between mice in the same groups, we observed no significant difference in bacterial translocation into the mLN between GF and SPF mice 22 hours after inoculum (Suppl.Fig.3B). Interestingly, *B.theta* CFU in the mesenteric lymph nodes were later reduced in all mice, with almost all SPF mice showing no counts in mLN at 72 hours post-inoculation (Suppl.Fig.3B).

In order to modulate the antigenic load in the gut, we generated a *B.theta* strain deleted for the BT4295 locus that encodes the epitope recognized by BΘOM T cells (KO strain). By mixing the wildtype (WT) and KO strains, we can control the abundance of cognate antigen, relatively independently of the total *B.theta* population density. We therefore orally inoculated GF mice with a ratio of 1:10 between the *B.theta* WT and KO strains (Low WT, Fig.1F). This resulted in a more than a 10-fold decrease in the abundance of the BΘOM T cell epitope carrying strain (Low WT) compared to monocolonized GF mice (Fig.1G). Surprisingly, although we still observed a high percentage of BΘOM T cells that divide and are activated in the Low WT group as compared to mock-colonized or KO-*B.theta*-monocolonized mice, this lower antigen dose fails to support T cell expansion suggesting that T cells are dividing but fail to survive (Fig.1H). Similarly, we increased WT *B.theta* luminal load in SPF mice by depleting the microbiota with broad-spectrum antibiotics before *B.theta* colonization (Fig.1I), obtaining *B.theta* gut loads almost 100-fold higher than in non-pre-treated SPF mice (Fig.1J). Nevertheless, despite the clear increase on BΘOM T cell division in the mice colonized with *B.theta* WT compared to the KO strain; transient microbiota depletion was not sufficient to support BΘOM T cell expansion (Fig.1K).

In conclusion, *B.theta* colonization can induce BΘOM T cell division and expansion of T cell numbers only at high bacterial antigen loads in GF mice. Titrating the antigenic load in the gut lumen of both GF and SPF mice generates expected changes in the observed level of BΘOM T cell division, but with unexpected effects on BΘOM T cell expansion, suggesting that suboptimal stimulation fails to generate sufficient survival signals for dividing T cells. This data also indicates that antigen load alone fails to explain the observed difference in T cell proliferation between SPF and GF mice.

### Gut pre-colonization increased the threshold for BΘOM T cell activation in immunocompetent mice

Another contributor to the difference in BΘOM T cell activation in GF and SPF mice could be that previous intestinal bacterial exposure alters the gut microenvironment to increase the threshold for T cell activation against new incoming bacteria. To test this idea, we pre-colonized GF mice using the *B.theta* KO strain together with the commensal *Eubacterium rectale* ATCC 33656 *(E.rectale)* and *E.coli* HS. Six days post initial colonization, the *B.theta* KO strain was replaced by an erythromycin-resistant *B.theta* WT strain by short-term supplementation of erythromycin into the drinking water (Col+WT group, Fig.2A). We included as controls a pre-colonized group of mice without *B.theta* KO (Col+KO group), and another group colonized only with *B.theta* WT as described before (WT group, Fig.1A). At the day of adoptive transfer, all groups colonized with *B.theta* WT strain had similar bacterial loads in the feces (Fig.2B).

**Figure 2.**
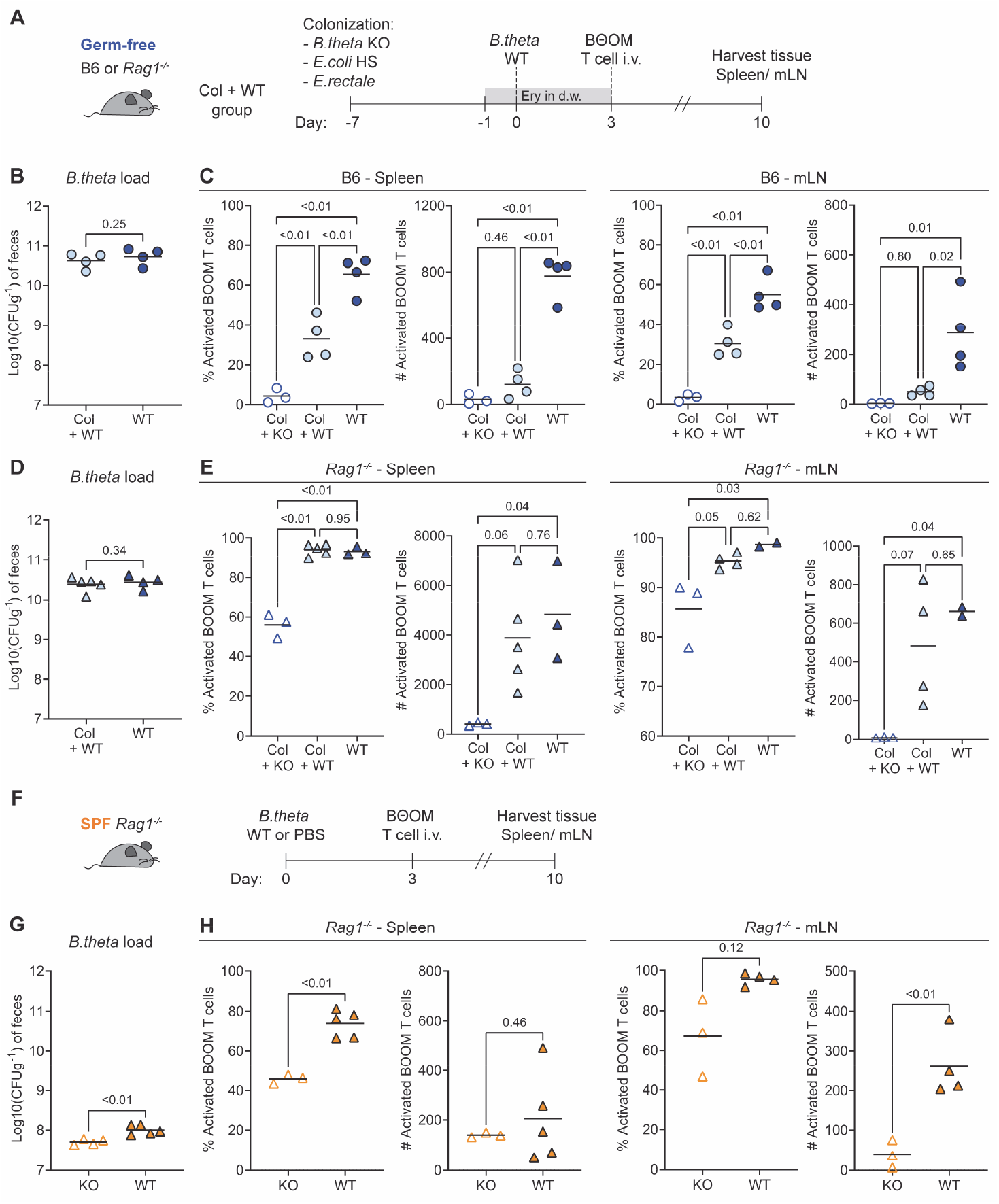
Gut colonization does not restrict BΘOM T cell activation in immunodeficient mice. **(A)** Experimental set-up of pre-colonization of GF wild-type (B6, circles) and immunodeficient *(Rag1^-/-^*, triangles) mice. Mice were initially colonized with *E.rectale, E.coli* HS and *B.theta* KO strains. One week after, mice received erythromycin in the drinking water and were colonized with an erythromycin-resistant *B.theta* strain (Col+WT). As controls, one group was kept undisturbed after the initial colonization till adoptive transfer (Col+KO) and another group was colonized only with *B.theta* WT as described before (WT, Fig.1A). **(B and D)** Gut luminal load of *B.theta* strains at the time of BΘOM T adoptive transfer in GF **(B)** B6 and **(D)** *Rag1^-/-^* mice. **(C and E)** Percentage and number of activated BΘOM T cells in spleen and mLN in GF **(C)** B6 and **(E)** *Rag1^-/-^* mice. **(F)** Experimental set-up of *B.theta* colonization in SPF *Rag1^-/-^* mice. **(G)** Gut luminal load of *B.theta* strains at the time of BΘOM T adoptive transfer **(H)** Percentage and number of activated BΘOM T cells in spleen and mLN.

Pre-colonization of GF mice with a simple microbial community significantly reduced the percentage and number of activated BΘOM T cells induced by *B.theta* WT in both spleen and mLN (Fig.2C). In fact, total numbers of activated BΘOM T cells in the pre-colonized group were undistinguishable from the control carrying the *B.theta* KO strain (Fig.2C). Interestingly, in both groups colonized with *B.theta* WT, we observed that approximately 40 to 60% of activated BΘOM T cells became Tregs in spleen and mLN (Suppl.Fig.4A).

As competition between lymphocytes plays a major role in regulating bacteria-host interactions in the gut [3]; we tested whether the absence of the adaptive immune system in *Rag1^-/-^* mice, influenced how pre-colonization affected activation of BΘOM T cells in the gut. We pre-colonized GF *Rag1^-/-^* mice with a 3-species microbiota as described above (Fig.2A) and observed similar *B.theta* loads as in WT GF mouse colonization at the day of adoptive transfer (Fig.2D). Contrary to what we observed in WT mice, pre-colonization did not reduce the activation of BΘOM T cells induced by *B.theta* WT colonization (Fig.2E) and most of activated BΘOM T cells did not acquire a regulatory phenotype (Suppl.Fig.4B) in *Rag1^-/-^* mice.

We also tested effect of immunodeficiencies in *bona fide* SPF colonized mice (Fig.2F), in which gut *B.theta* loads are more restricted (Fig.2G). Although we observed a tendency towards higher proliferation of BΘOM T cells in *Rag1^-/-^* SPF mice colonized with *B.theta* WT compared to the control KO strain, there was no major increase in the cell numbers as observed in GF *Rag1^-/-^* mice (Fig.2H) and T cells did not acquire a regulatory phenotype (Suppl.Fig.4C). However, it should be pointed out that the interpretation of these data is complicated by potential cross-reactivity of BΘOM T cells to epitopes in the endogenous SPF microbiota. Finally, we probed the effect of acute depletion of Tregs on *B.theta*-specific T cell activation in fully colonized conditions using DEREG SPF mice (Suppl.Fig.5A). After depletion of host Tregs (Suppl.Fig.5B), BΘOM T cells were activated regardless of the *B.theta* strain used for colonization (Suppl.Fig.5C), indicating antigen-independent proliferation or activation by other cross-reactive microbiota members.

In summary, despite comparable *B.theta* antigen load in the gut, there is a differential impact on BΘOM T cell proliferation depending on the integrity of the gut immune system. In the absence of endogenous T and B cells, pre-exposure to gut microbes no longer prevents specific T cell activation, indicating an important role for other lymphocytes in controlling the threshold for intestinal T cell activation. However, Treg depletion resulted in strong antigen-independent T cell proliferation suggesting that Treg activity alone is not sufficient to explain this observation.

### T cell activation by gut bacteria depends on the TCR-transgenic model

TCR transgenic models are well known to show TCR-specific biases in experimental outcome. We therefore sought to confirm our observations in a second TCR transgenic system. To achieve this, we designed a plasmid that allows for surface display of an inserted peptide sequence in the non-pathogenic *E.coli* BL21 strain. We generated two different plasmids to allow for display of the BΘOM T cell Ag (BT4295_541-554_, *E.coli*-BT425) or the SMARTA-1 T cell (SM-1) Ag (GP_64-80_ from lymphocytic choriomeningitis virus (LCMV), *E.coli-GP64)* (Suppl.Fig.6A). Both *E.coli* strains expressed similar levels of surface epitopes (Suppl.Fig.6B) and when administered systemically (Suppl.Fig.7A) were able to activate BΘOM (Suppl.Fig.7B) and SM-1 T cells (Suppl.Fig.7C).

We then tested the capacity of these strains to activate BΘOM and SM-1 T cells during gut colonization. After inoculating GF B6 or *Rag1^-/-^* mice with either *E.coli*-BT425 or *E.coli*-GP64, we transferred BΘOM and SM-1 labelled T cells at a 1:1 ratio (Fig.3A and Suppl.Fig.8). Bacterial loads of both *E.coli*-BT4295 and *E.coli*-GP64 were similar in both B6 (Suppl.Fig.9A) and *Rag1^-/-^* mice (Suppl.Fig.9B); although around 100-fold lower than the usual *B.theta* CFU density in GF mice (Suppl.Fig.3A).

**Figure 3.**
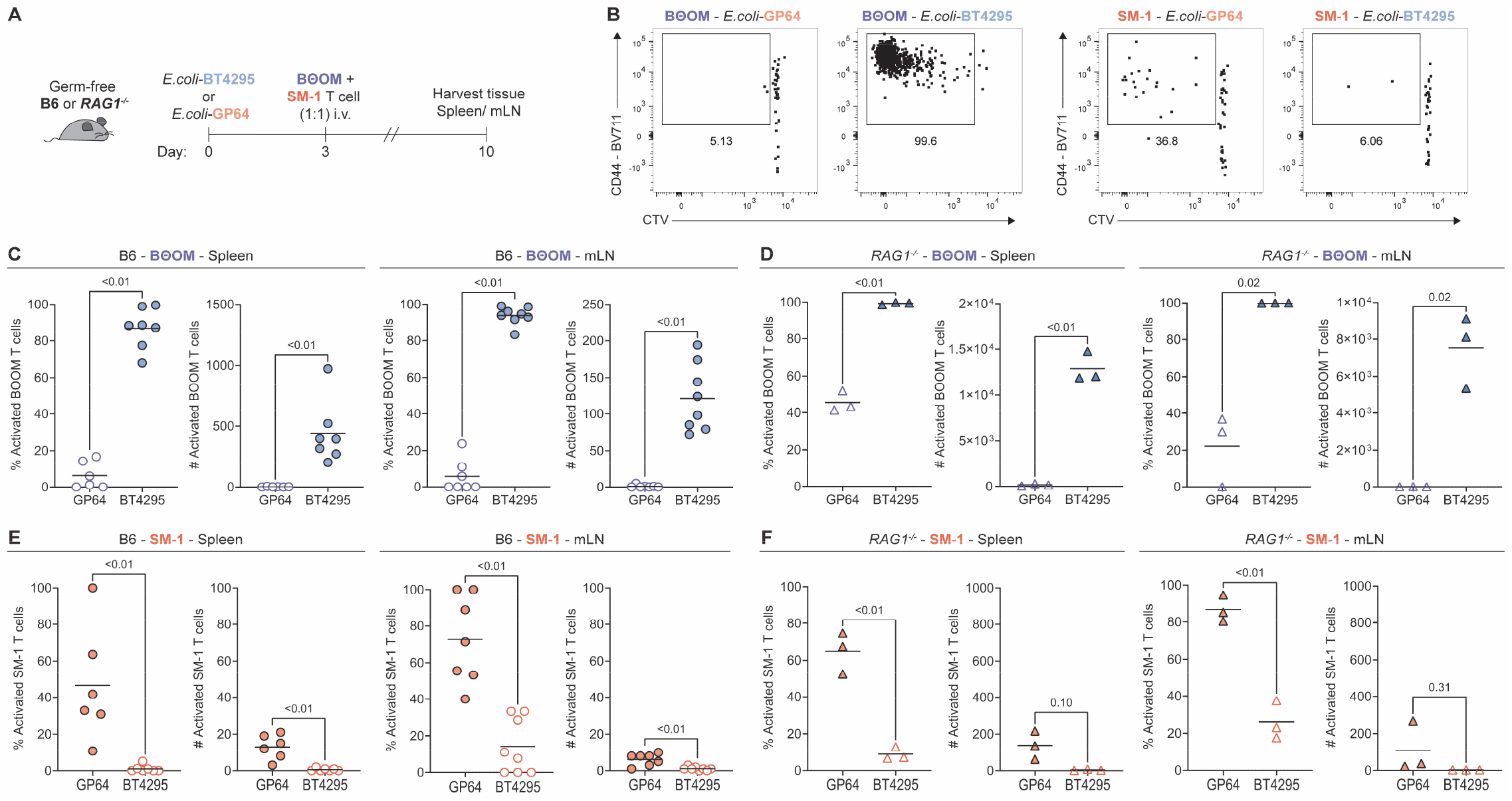
BΘOM and SM-1 T cell proliferate differently to gut colonization with antigen-expressing bacteria. (A) Experimental set-up of C57BL/6J (B6) or *Rag1-/-* GF mice colonization with antigen-expressing *E.coli* BL21. Strains carry a plasmid for surface display of either the *B.theta* BT429_541-554_ (*E.coli*-BT4295) or the LCMV GP_64-80_ (*E.coli*-GP64) epitope. BΘOM T cells and SM-1 T cells were adoptively transferred in a 1:1 ratio. (B) Representative flow cytometry plot of activated BΘOM and SM-1 T cells. (C and D) Percentage and number of activated BΘOM T cells in spleen and mLN in GF (C) B6 and (D) *Rag1^-/-^* mice. (E and F) Percentage and number of activated SM-1 T cells in spleen and mLN in GF (E) B6 and (F) *Rag1^-/-^* mice.

Despite the lower CFU densities, BΘOM T cells proliferated in *E.coli*-BT4295 colonized B6 mice (Fig.3B), reaching similar levels of activated T cells (Fig.3C) and Tregs (Suppl.Fig.9B) as those observed during WT *B.theta* colonization (Fig.2B, Suppl.Fig.4B). BΘOM T cell proliferation was more noticeable in GF *Rag1^-/-^* mice. We found a massive expansion in the total number of BΘOM T cells in *E.*coli-BT4295 colonized ex-GF *Rag1^-/-^* mice (Fig.3D), which was 4-10 times more than the one observed during *B.theta* colonization (Fig.2C). As previously observed during *B.theta* colonization (Suppl.Fig.4D), almost no BΘOM T cells acquired a regulatory phenotype in *E.*coli-BT4295 colonized ex-GF *Rag1-/-* mice (Suppl.Fig.9C).

On the other hand, SM-1 T cells in GF B6 mice colonized with the corresponding *E.coli*-GP64 strain did not expand (Fig.3B). Although there was cell division (Fig.3E), there was no comparable increase in cell numbers (Fig.3E). In the case of GF *Rag1^-/-^* mice colonization with *E.coli*-GP64, SM-1 showed antigen-specific expansion with almost no homeostatic proliferation in control *E.coli*-BT4295 colonized mice (Fig.3F). Interestingly, the few activated SM-1 T cells differentiated into Tregs in a similar proportion as activated BΘOM T cells (Suppl.Fig.9D and Suppl.Fig.9E). Based on this data, either antigen presentation of the SM-1 peptide epitope or activation of SM-1 cells in the gut is dramatically less efficient than BOOM T cells.

## DISCUSSION

Bacterial antigens are constantly interacting with the immune system in the gut, and many of the adaptive immune cells in the gut are microbiota specific during homeostasis[11],[12]. We explored some fundamental aspects of this interaction by studying the role of gut colonization and bacterial load in regulating the activation threshold for T cells in the gut.

BΘOM T cell were activated and proliferated in ex-GF B6 mice after colonization with *B.theta,* similarly to previously reported with antibiotic-treated SPF *Rag1-/-* mice[10]. However, we observed that SPF B6 mice restricted BΘOM T cell proliferation after colonization with *B.theta.* It should be noted that *B.theta* can establish in the gut of our SPF mice in a stable fashion[13]. Also, this restriction does not seem to be mediated by an intrinsic difference in the immune system of GF and SPF mice as adoptively transferred BOOM T cells respond similarly to a systemic challenge with inactivated bacteria in both environments. Similarly, *B.theta* sampling into the mLNs appeared to be independent on the colonization state at early time-points and showed similar loads as other non-invasive commensal bacteria[14],[15].

One potential mechanism that restricts BΘOM T cell activation is the bacterial antigen load in the gut, which is determined by the luminal bacterial density and the antigen expression per bacterial cell. In the case of *B.theta,* high bacterial density is required to induce BΘOM T cell proliferation in GF mice. Although lower densities of *B.theta* failed to induce substantial proliferation in GF mice, this can be related to the level of antigen expression per cell. Reducing gut bacterial density by diluting the antigen-expressing *B.theta* with the KO strain, which competes for a very similar niche in the gut, diminishes the number of antigen-producing bacteria that are sampled into the mLN during the initial hours post-colonization (1-10 bacteria instead of 1000). Interestingly, *E.coli* expressing the BT4295 peptide was equally stimulatory to BOOM T cells in GF mice despite colonizing to around a 100-fold lower level. This could be due to over-expression of the antigen compared to endogenous levels in *B.theta,* more efficient antigen processing than the endogenous *B.theta* protein, or increased adjuvanticity of the *E.coli* presenting the antigen.

In the case of SPF mice, when antibiotic pre-treatment was used to colonize *B.theta* to levels close to those observed in GF mice, we could measure a small amount of BΘOM T cell activation. However, T cell proliferation and expansion remained very limited, which indicates mechanisms operating above and beyond the gut antigen load. We hypothesized therefore that BΘOM T cell activation and proliferation could be restricted in SPF mice by the immune modulation imprinted by their microbiota. Indeed, bacterial colonization in GF mice, either by a single strain[16] or a consortium[17], can modify responsiveness of the gut immune system. In addition, it has been shown that microbiota regulation of innate immunity can affect the proliferative capacity of transferred T cells[18]. Consistent with this, pre-colonizing mice with a very simple community, which includes representatives of the main gut phyla, damped the activation and proliferation levels of BΘOM T cell. However, the mechanism behind this regulation seems to be related to a functional adaptive immune system, as no such downregulation was observed in *Rag1^-/-^* animals. Depletion of Tregs induced an entirely different phenomena, also suggesting that the regulation may be based on cell-cell competition rather than insufficient Treg activity per-se in GF mice. Therefore, it seems that after initial colonization by bacteria, the adaptive immune system sets the microenvironment of the gut to a higher threshold for activation, preventing newly-incoming from bacteria inducing major activation and proliferation of T cells.

Finally, there are some limitations in the use of TCR-Tg cells for studying antigen specific responses. Different TCR-Tg T cells potentially behave different under similar antigen exposure based on their antigenic affinity or the surrounding microenvironment. For example, *Akkermansia-specific* TCR-Tg T cells are able to proliferate in colonized mice, both gnotobiotic and SPF[19]. Compared to BΘOM T cells, we observed a much weaker activation of SM-1 T cells, with almost no proliferation in monocolonized B6 mice and only a very limited expansion in *Rag1-/-.* This limited expansion of TCR-Tg T cells has been reported in a similar model using *E.coli* for antigen delivery, even under barrier disrupting conditions such DSS treatment[20],[21]. Assuming that antigenic exposure in the gut is similar, we should ponder other potential mechanisms by which we and others found this restricted proliferation of SM-1 T cells. For example, we found that SM-1 have an approx. 200 times higher affinity (EC50 ~5nM,[22]) towards their cognate peptide than BΘOM T cells (Suppl.Fig.10). Therefore there is no simple linear relationship between T cell activation and affinity. However, we cannot exclude that this plays a role, for example via stronger interactions with tolerance mechanisms such as clonal deletion[23]–[25]. CBir1, another microbiota-specific TCR-Tg T cells, are also unable to proliferate in the gut of immunocompentent B6 mice[26], as they are deleted in an antigen-dependent manner upon activation[27]. Exploring the basic mechanisms that influence why different TCR-expressing cells undergo clonal deletion or phenotypic differentiation in the gut and their similarities to affinity-based selection in the thymus will be an exciting topic of future studies.

In summary, we found that the activation threshold of gut T cells towards incoming bacteria is strongly influenced by pre-exposure to bacteria in a manner dependent on an intact adaptive immune system. High antigen loads, and the immunostimulatory nature of the antigen-expressing bacteria contribute to generating robust T cell activation in bacteria-naïve mice, but fail to induce strong T cell proliferation of BOOM T cells in pre-colonized gnotobiotic or fully colonized mice.

## ABBREVIATIONS

B6: C57BL/6 mice
*B.theta*: *Bacterioides thetaiotaomicron*
BΘOM: T-cell receptor transgenic specific for *B.theta* BT4295 epitope
CFU: colony-forming units
CTV: CellTrace Violet
GF: Germ-free
LCMV: Lymphocytic choriomeningitis virus
mLN: mesenteric lymph nodes
*Rag1^-/-^*: RAG1 knock-out mice
SM-1: T-cell receptor transgenic specific for LCMV BT4295 epitope
SPF: Specific-pathogen free
TCR-Tg: T-cell receptor transgenic
Treg: regulatory T cell

## ACKNOWLEDGEMENTS

We would like to thank Prof. Paul Allen for kindly sharing the BΘOM TCR-Tg mouse line with us. We would also like to express our gratitude Dr. Roman Spörri and Prof. Annette Oxenius for providing SMARTA-1 TCR-Tg mice, experimental reagents and for providing a discussion environment for this project in their group meetings. Prof. Eric Martens kindly supplied the parental *B.theta* strain and the pNBU2 plasmid carrying a tetracyclin resistant cassette (TetQ). Finally, we thank Prof. Carolyn King, Dr. Daniela Latorre, Dr. Ilaria Spadafora, Dr. Ioana Sandu and Verena Lentsch for enjoyable discussions and the interesting ideas about this project.

## Author Contributions

Conceptualization, D.H., and E.S.; Methodology, D.H., A.E., T.K., N.J. and E.S.; Formal Analysis, D.H.; Investigation, D.H., B.C., A.E., S.B., E.F., C.M.; Resources, T.K., N.J., and E.S.; Writing – Original draft, D.H. and E.S.; Writing – Review and Editing D.H., B.C., A.E., S.B., E.F., C.M., T.K., N.J. and E.S.; Visualization, D.H., and E.S.; Supervision, T.K., N.J. and E.S.; Funding acquisition, N.J. and E.S.

## METHODS

### Mice

GF B6 mice are bred and maintained in open-top cages within flexible-film isolators, supplied with HEPA filter air, and autoclaved food and water. GF status of these colonies is monthly assessed via anaerobic and aerobic liquid cultures. SPF B6 mice are bred and maintained in IVC cages in a clean mouse facility. All mice used in the experiments were adults between 12-18 weeks old, males and females. *Rag1^-/-^* mice [28] were rederived under GF conditions. *Rag1^-/-^* mice GF and SPF mice were kept under conditions as described before at the ETH Phenomics Center (EPIC). DEREG mice [29] were kept under SPF conditions and experiment were performed at the Laboratory Animal Services Center (LASC) at the University of Zürich.

Sperm from a BΘOM CD45.1^+/+^ *Rag1^-/-^* transgenic mouse line was rederived at EPIC [10]. BΘOM CD45.1+/− *Rag1-/-* transgenic mice are kept in a clean mouse facility under enhanced SPF conditions. SMARTA (SM-1) CD45.1 +/−transgenic mice are bred and maintain under similar conditions[30]. All experiments were conducted in accordance with the ethical approval of the Zürich Cantonal Authority under the license ZH120/19.

### Bacterial strains

All *Bacteroides thetaiotaomicron (B.theta)* strains were produced by using *Δtdk* VPI-5842 strain as background *(B.theta* WT strain). The KO strain *(BT4295* gene deletion) was produced via counterselectable allelic exchange as described before [31]. Briefly, a *Δtdk* strain conjugated with the pExchange-tdk-ermG vector, carrying the 1Kb flanking homologous regions of the *BT4295* gen. After initial homologous recombinantion, clones underwent selection in brain-heart infusion (BHI) agar plates supplemented with of 10% sheep blood (BHI-blood) plates with erythromycin (25 μg/ml) plus gentamycin (200 μg/ml). Isolated clones were counterselected in BHI-blood plates with 5-fluoro-2’-deoxyuridine (200 μg/ml) which forces a second homologous recombination. Selected clones were screened by PCR and *BT4295* deletion was confirmed by sequencing.

In both strains, *B.theta* WT and KO, we introduced a short-genetic tag, a fluorescent protein (GFP or mCherry) and antibiotic resistance (against erythromycin or tetracycline) by using the mobilizable Bacteroides element NBU2, which integrates into the Bacteroides genomes at a conserved location [32]. The suicide NBU2 plasmid carrying the described inserted genes was transferred to the target *B.theta* strains by conjugation with *E.coli* S17-1. *B.theta* strains that integrate the suicide NBU2 plasmid were selected in BHI-blood plates supplemented with gentamycin (200μg/ml) and either erythromycin (25μg/mL) or tetracycline (2μg/mL) depending on the antibiotic resistance transferred in the plasmid. After 48 hours, single colonies were streaked in fresh BHI-blood agar plates with antibiotics, to avoid potential contamination with WT strains. Successful insertion in the BTt70 or BTt71 sites was evaluated by PCR.

For *E.coli* strains, constitutive expression of GP64 and BT4295 T-cell peptides on the surface was achieved by transformation with the plasmids pTK358 and pTK557, respectively, and maintenance in selective media. The plasmids consisted of a kanamycin resistance gene, an RSF1030 origin of replication, an expression cassette driven by the weak constitutive promoter J23114 [33], and an open reading frame encoding a PelB secretion signal, C-Myc-tag, the T cell peptide, followed by the neck, stalk and transmembrane domain (aa973-1098) of the trimeric autotransporter adhesin Hia from *H. influenzae* [34]. Surface display of the peptide was assessed by bacterial flow cytometry [35] using anti-Myc-AF647 (9B11) antibody.

### Bacterial cultures

*B.theta, Eubacterium rectale* ATCC 33656 *(E.rectale)* and *E.coli* HS strains were streaked from frozen stocks on brain-heart infusion (BHI) agar plates supplemented with of 10% sheep blood (BHI-blood agar) and grown anaerobically (5% H_2_, 10% CO_2_, rest N_2_) at 37°C for at least 48 hours. Similarly plasmid carrying *E.coli* BL21 strains were streaked on LB plates supplemented with kanamycin (50μg/mL) and grown aerobically at 37°C overnight.

For the preparation of inoculums for colonization, several colonies were picked and grown anaerobically in standing cultures of brain-heart infusion supplemented media (BHIS: 37 g/L BHI (Sigma); 1 g/L-cysteine (Sigma); 1 mg/L Hemin (Sigma)) at 37°C for 12-18 hours. Liquid cultures were supplemented with either erythromycin (25 μg/mL) or tetracycline (2 μg/mL) depending on the antibiotic resistance inserted in the strain. In the case of plasmid carrying *E.coli* BL21 strains, single colonies were picked and grown aerobicaly, in shaking cultures of LB supplemented with kanamycin (50 μg/mL) at 37°C for 12-18h.

### Bacterial inoculums for colonization

For colonization experiments in GF, gnotobiotic and SPF mice, we used strains described on the specific figures. For all colonization experiments, all cultures were spun down (3000rcf for 20min at 10°C) and washed once with cold PBS buffer to eliminate any antibiotic from the cell suspention. Bacterial density was quantified via optical density (1 O.D. ~ 4×10^8^ bacteria/mL) and bacterial numbers were adjusted to approximately 10^8^ bacteria/mL. Usual dose of colonization for all experiments was approximately 10^7^ bacteria, unless otherwise specified in the experimental setting, delivered by gavaging 100 μL.

For experiments in Fig.1 comparing different ratios of erythromycin-resistant WT and tetracyclin-resistant KO *B.theta* strains, we kept the control *B.theta* KO strain in 10^7^ bacteria per inoculum and added 10^5^ of *B.theta* WT to the inoculum. For experiments in Fig.2 assessing the effects of pre-colonization, *B.theta* KO, *E.rectale* and *E.coli* HS were grown in individual liquid cultures as described before. While in the anaerobic tent, cultures were mixed in a 1:1 ratio depending on the experimental group *(B.theta* KO + *E.rectale* + *E.coli* HS or *E.rectale* + *E.coli* HS) and sealed in a sterile and anaerobic serum bottle. Cultures were under anaerobic conditions until some minutes before being gavaged.

### Bacterial load quantification

Colonization status and bacterial loads of all strains were assessed by culture in agar plates. Fecal pellets, cecal content or mesenteric lymph nodes (mLN) were sampled at the designated timepoints and weighed (except mLNs). All samples were homogenized in PBS (1 mL for cecal content and 0.5 mL for fecal pellets and mLNs) for 2.5 min at 25 Hz in a TissueLyser (Qiagen, Germany). In the case of mLNs, 200 μL of the homogenized tissue was plated on BHI-blood agar plates supplemented with erythromycin (25 μg/mL). For fecal and cecum samples, a serial dilution was performed in PBS and 10ul of each dilution was plated as parallel lanes in agar plates. For experiments including *B.theta,* we plated the serial dilution of fecal or cecum samples in BHI-blood agar plates supplemented with either erythromycin (25 μg/mL) or tetracycline (2μg/mL) depending of the strain used. Plates were incubated anaerobically at 37°C for at least 48 hours. For experiments including *E.coli* strains, we plated the serial dilution of fecal samples in LB agar plates supplemented with kanamycin (50 μg/mL), and then incubated aerobically at 37°C overnight. Colonization status and bacterial loads of *E.rectale* and *E.coli* HS strains were assessed by qPCR.

### Adoptive T cell transfers

BΘOM T cells were obtained from TCR-Tg+/− CD45.1+/− *Rag1-/-* mice. SMARTA-1 (SM-1) T cells were obtained from TCR-Tg+/− CD45.1+/− *Rag1+/−* mice. Donor mice matched the gender of the reciepient mice whenever possible, otherwise female donors were used. Spleen was harvested and disintegrated into single suspension using a 40μm cell strainer (Falcon) and MACS buffer (PBS, 2% FBS, 5mM EDTA). After washing cells once (800 rcf, 5 min, 4°C), we positively selected CD4+ T cells using CD4 MicroBeads and magnetic sorting (Miltenyi Biotec). Enriched CD4+ T cells were counted, washed in PBS, adjusted to 2×10^6^ cells/mL concentration in PBS and with 5μM of CellTrace™ Violet for 20min at 37°C in water bath. After quenching the reaction with 5 volumes MACS buffer, cells were washed again in PBS and adjusted to a concentration of 1×10^6^ cells/mL. Mice were injected with 2×10^5^ CTV-labelled T cells in a volume of 200 μL via the tail vein.

### Acute regulatory T cell depletion

DEREG and C57B6/L mice were injected with 200 ng of diphtheria toxin (DT, Merck) one day before adoptive T cell transfer, and one and three days afterwards. Total body mass was monitored pre-treatment and during the days of DT injection. Effect of DT on the depletion of regulatory T cells was confirmed at the end of the experiment by flow cytometry.

### Cell isolation from peripheral tissue

Spleen and mLNs were harvested 7 days after adoptive T cell transfer. mLNs were digested with 1U/mL of Liberase TL (Roche) and 50U/mL of DNAse I (Sigma-Aldrich) for 20 min at 37°C. Spleens and digested mLNs were disintegrated using a 40μm cell strainer (Falcon) into single cell suspension in MACS buffer (PBS, 2% FBS, 5mM EDTA). After washing the cells once in MACS buffer, we lysed red blood cells using RBC Lysis Buffer for 5min at RT (BioLegend). RBC Lysis Buffer was quenched with MACS buffer and washed once. Then, we enriched for CD4+ T cells by negative selection using biotinylated anti-CD8, anti-Ly-6G and anti-CD45R/B220 antibodies and MojoSort™ Streptavidin Nanobeads (BioLegend). In order to maximize cell recovery from spleen samples, we magnetically sorted the samples twice. After CD4 enrichment, cells were kept in ice until surface/intranuclear staining.

### Flow cytometry

For cell surface marker staining, the cell pellet was re-suspended in 100 μL of antibodies and LIVE/DEAD™ Fixable Near-IR dye (ThermoFisher Scientific) diluted in MACS buffer, and stained on ice for 30min. After washing once with MACS buffer, cells were fixed, permeabilized and stained for intranuclear transcription factors using the eBioscience™ Foxp3/Transcription Factor Staining Buffer Set acording to manufacturer indications (ThermoFisher Scientific). The following antibodies were used for surface and intranuclear staining: CD44-FITC (IM7, BioLegend), CD44-BV711 (IM7, BioLegend), TCR Vβ12-PE (MR11-1, BioLegend), TCR Vα2-PE/Cy7 (B20.1, BioLegend), CD45.2-PerCP (104, BioLegend), CD45.1-APC (A20, BioLegend), CD45.1-PE (A20, BioLegend), CD4-PE/Cy7 (RM4-5, BioLegend), CD4-PerCP (RM4-5, BioLegend), CD62L-FITC (MEL-14, BioLegend), CD62L-BV605 (MEL-14, BioLegend), Foxp3-FITC (FJK-16s, eBioscience), CD69-PE (H1.2F3,BioLegend).

### CD69 Activation Assay

BT4295 peptide in powder form were dissolved in RPMI-1640 (Gibco) and serial dilutions of the peptide were plated so that a dose-response curve could be generated. BΘOM T cells were isolated from total splenocytes of *Rag1^-/-^* CD45.1/2 BΘOM; and splenocytes were isolated from CD45.2/2 C57BL/6 mice as described before. We plated 10000 CD45.1/2 BΘOM T cells and 50000 CD45.2/2 splenocytes per well in RPMI-1640 supplemented with 10% FCS, 2 mM L-glutamine, 1% penicillin-streptomycin mix (50K U Pen/50 mg Strep), 1 mM sodium pyruvate, 0.1 nM non-essential amino acids, 20 mM HEPES and 50nM β-mercaptoethanol were added to the dilution series. Cells were incubated overnight at 37°C, and subsequently stained for CD69 surface expression.

### Statistics

We evaluated differences between two groups using Welch t test to assume for unequal standard deviation. For comparisons between more than two groups we used analysis of variance (ANOVA) with Tukey’s multiple comparisons test. All statistical tests were performed using the GraphPad Prism 9 software. P values of less than 0.05 were considered to be significant. In all graphs, we plotted individual values and the group mean.

## SUPPORTING INFORMATION

**Supplementary Figure 1:**
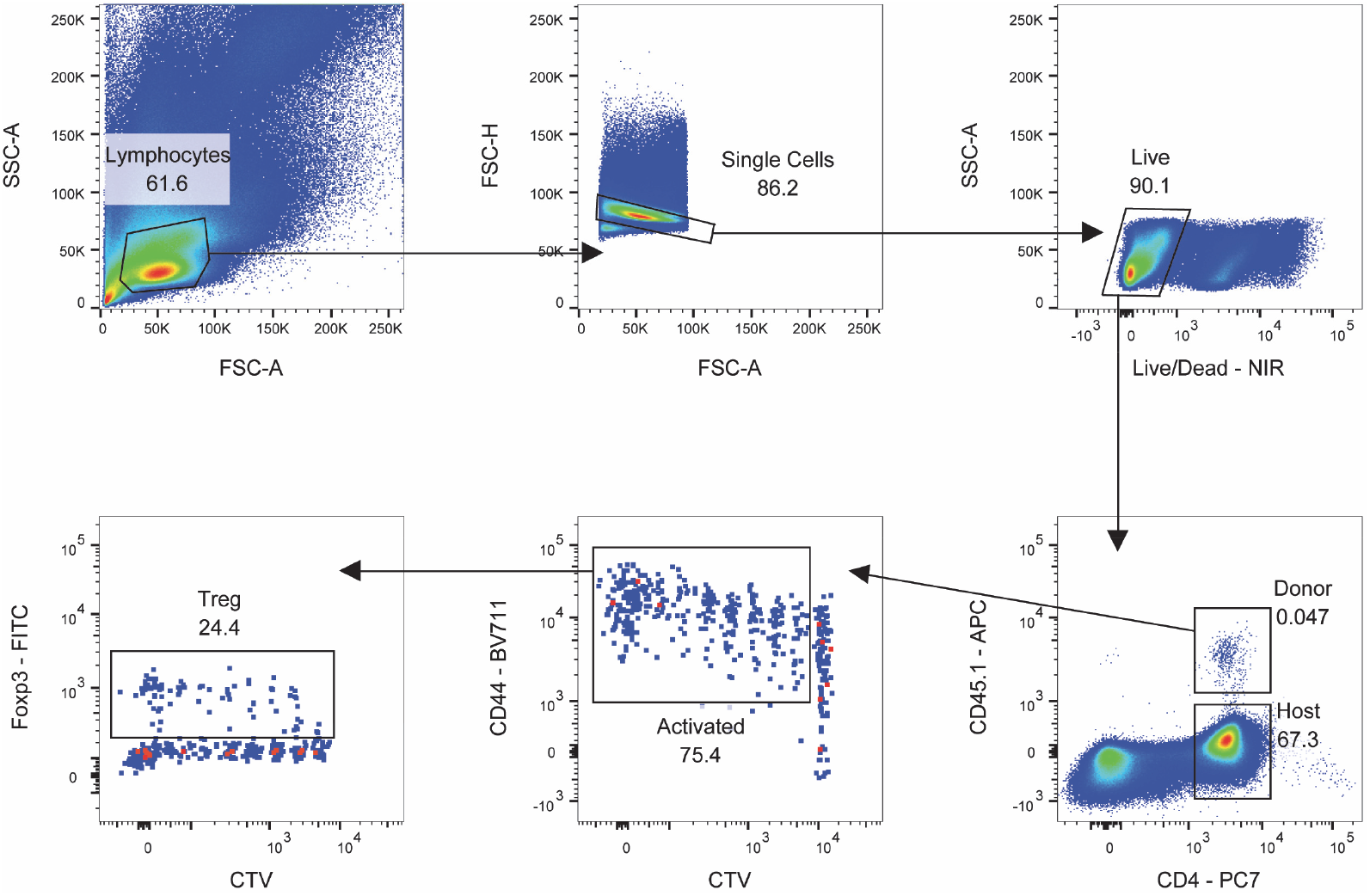
Gating strategy. After gating for live single-cell lymphocytes, donor BΘOM CD4+ T cells were identified by the expression CD45.1. Activated BΘOM T cells were gated by their expression of CD44 and dilution of the CTV proliferation dye. Controls as showed in Fig.1B were used to define the diluted and undiluted populations for the CTV dye.

**Supplementary Figure 2:**
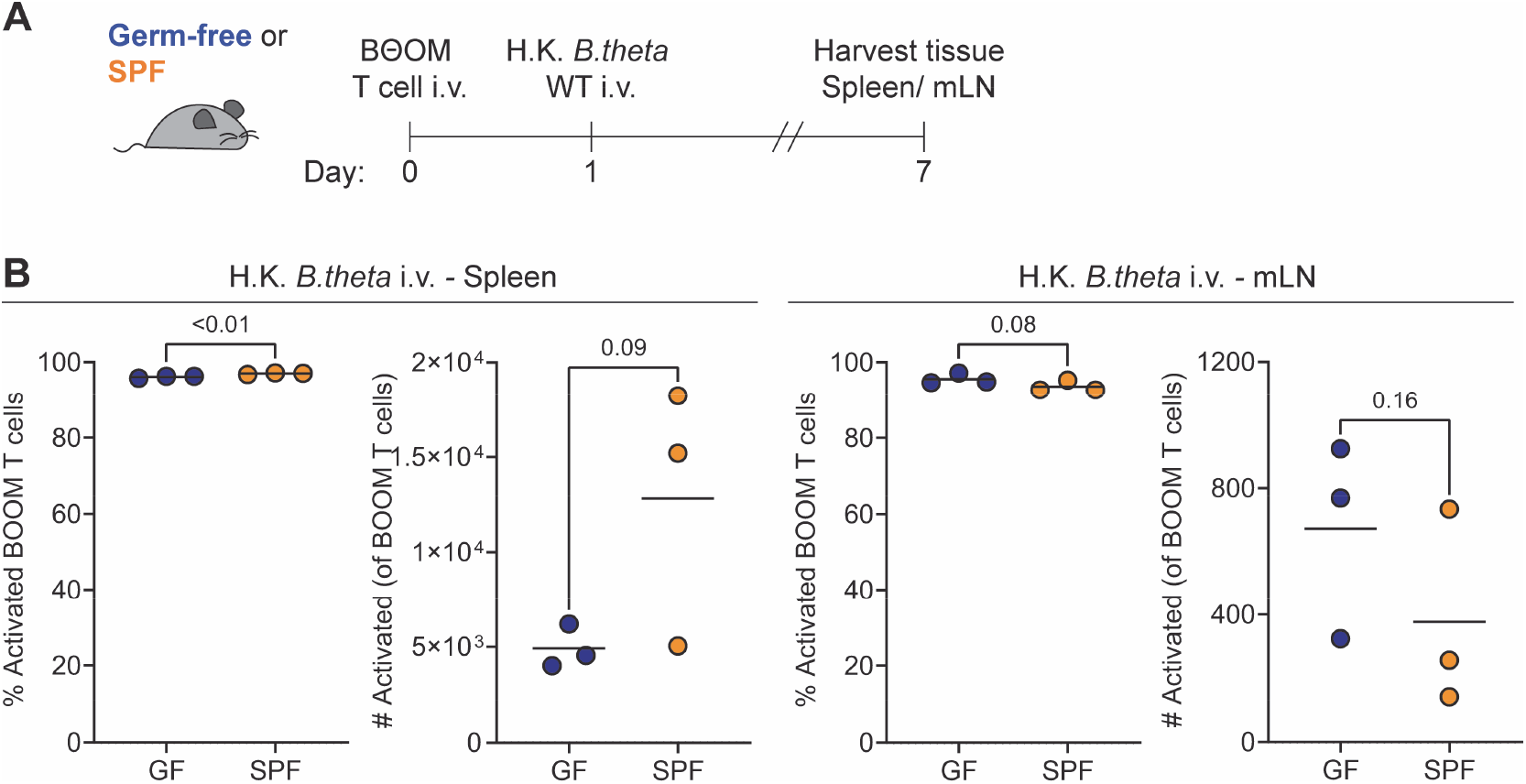
*B.theta* induce similar activation of BΘOM T cells by systemic challenge in GF and SPF mice. **(A)** Experimental set-up of *B.theta* systemic challenge. **(B and C)** Percentage and number of activated BΘOM T cells in GF and SPF mice both in **(B)** spleen and **(C)** mLN.

**Supplementary Figure 3:**
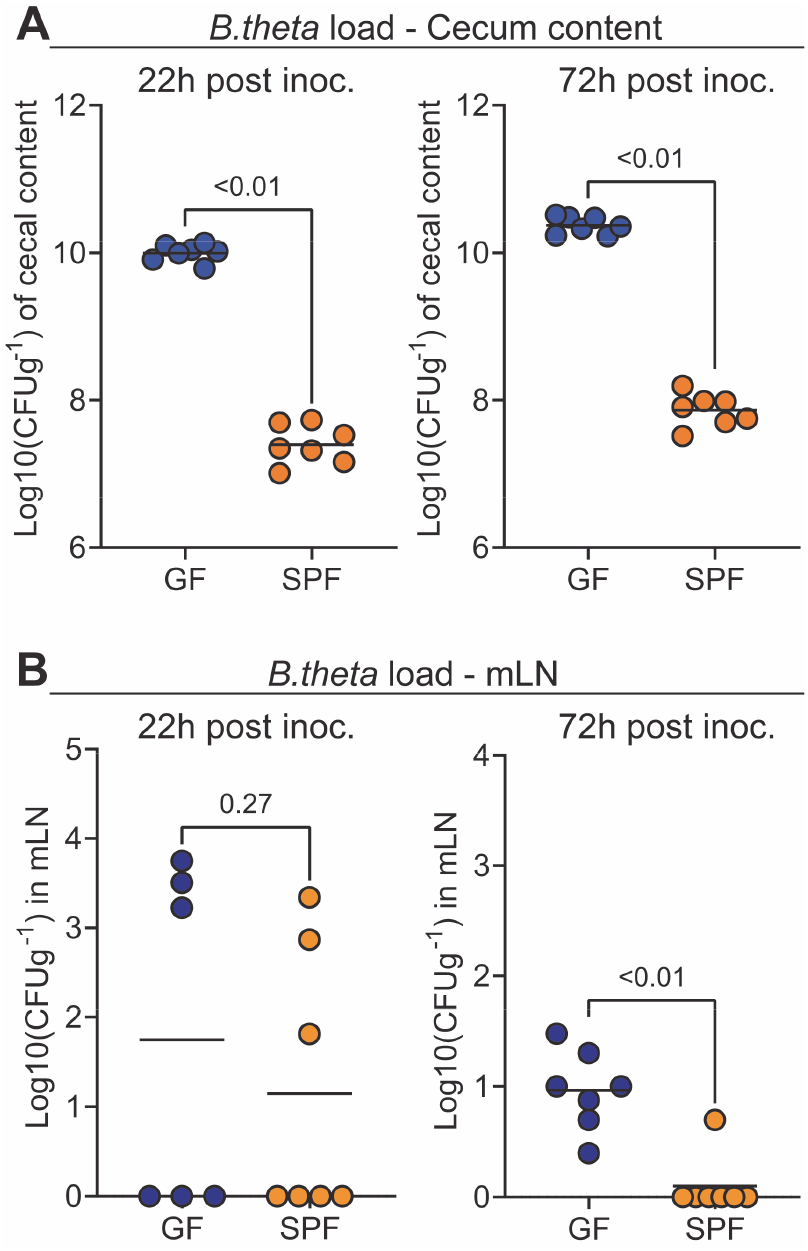
Bacterial load in gut lumen and mLN. **(A and B)** *B.theta* CFUs recovered from **(A)** cecum content and **(B)** mLNs of GF and SPF mice after 22 and 72 hours post oral inoculation.

**Supplementary Figure 4:**
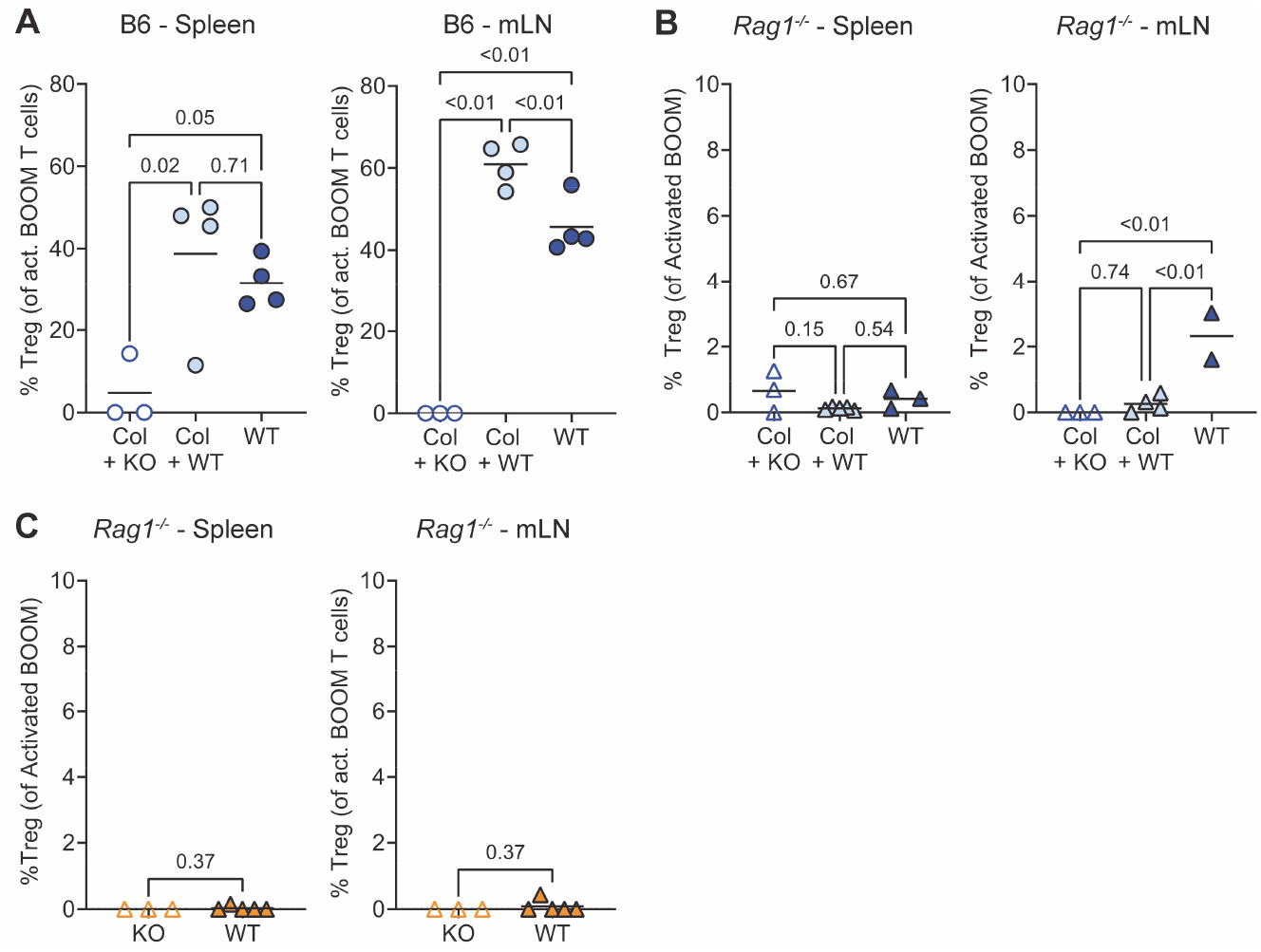
Bacterial load in gut lumen and induced BΘOM regulatory T cells. **(A-C)** Percentage of regulatory T cells (Treg) among activated BΘOM T cells in spleen and mLN. **(A)** Related to experiments described in Fig.2B-C, **(B)** Related to experiments described in Fig.2D-E. **(C)** Related to experiments described in Fig.2G-H.

**Supplementary Figure 5:**
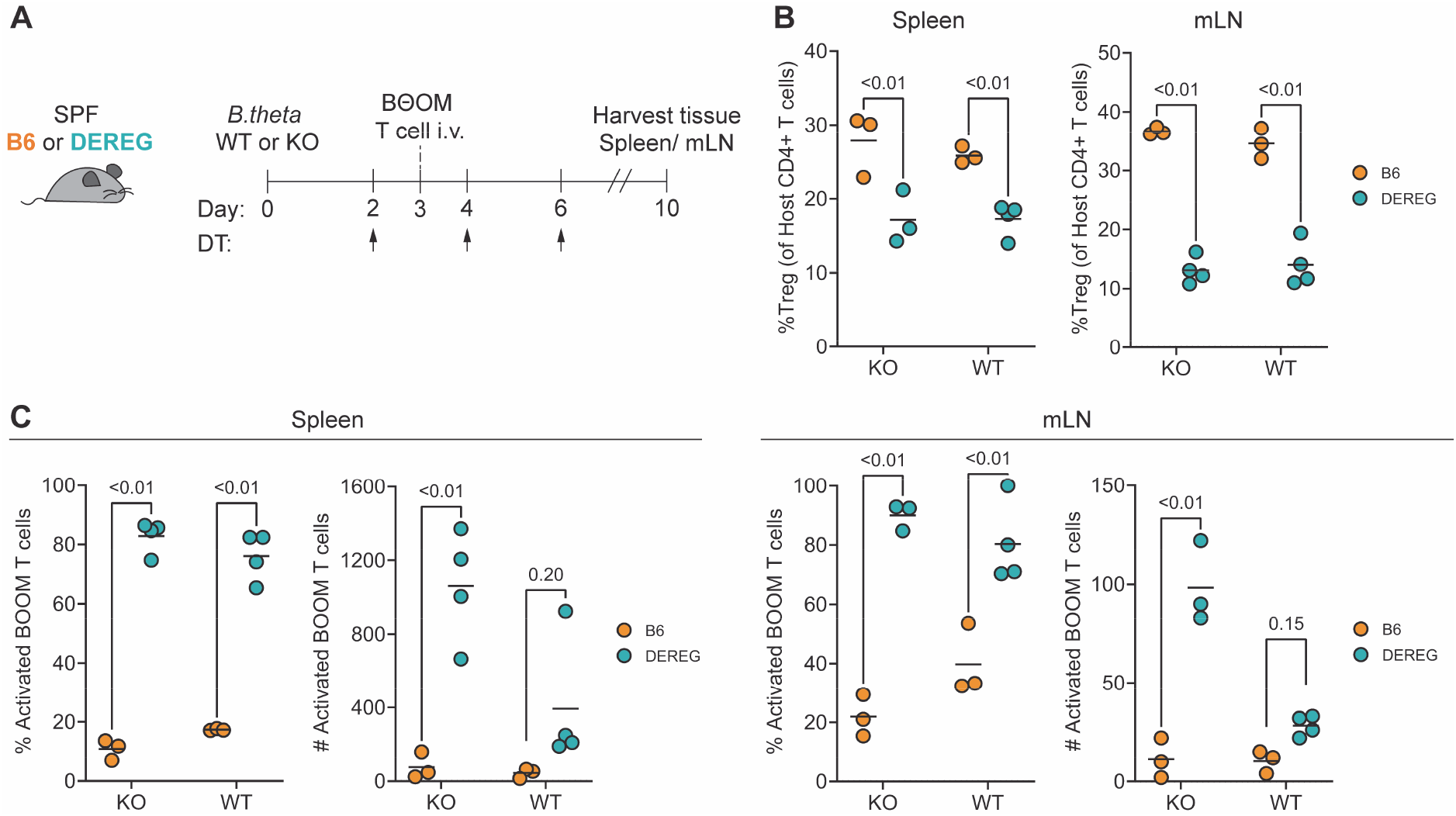
Acute depletion of regulatory T cells (Tregs). **(A)** Experimental set-up for acute regulatory T cell (Treg) depletion in DEREG mice. **(B)** Percentage of host Tregs among CD4+T cells at the end of the experiment in spleen and mLN. **(C)** Percentage and number of activated BΘOM T cells in B6 and DEREG mice both in spleen and mLN.

**Supplementary Figure 6:**
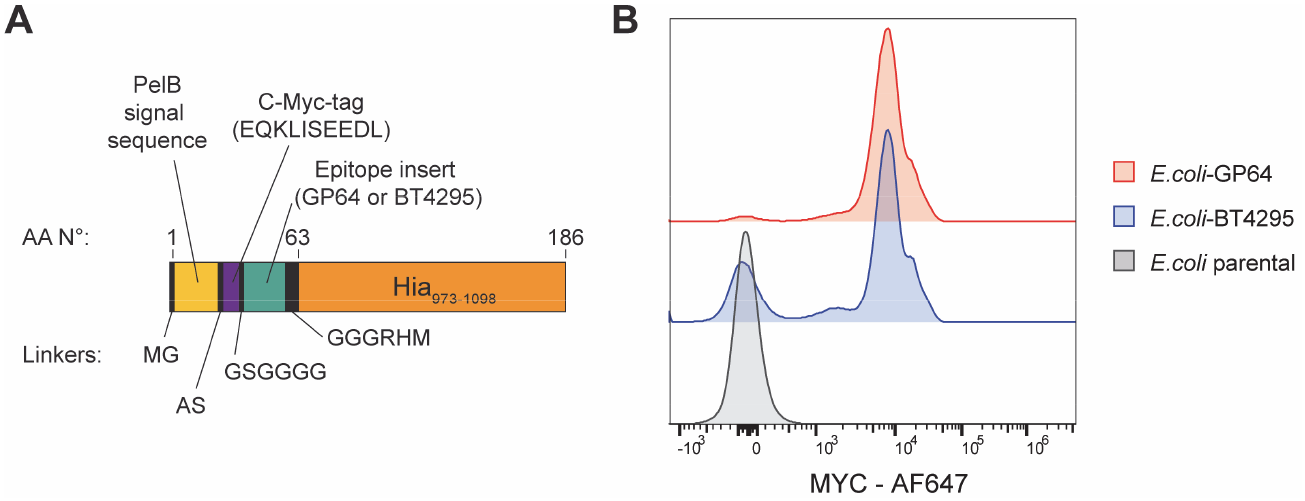
**(A)** Schematic representation of plasmid insert construct for expressing surface epitopes. **(B)** Representative flow cytometry plots depicted surface expression of epitopes in *E.coli* strains.

**Supplementary Figure 7:**
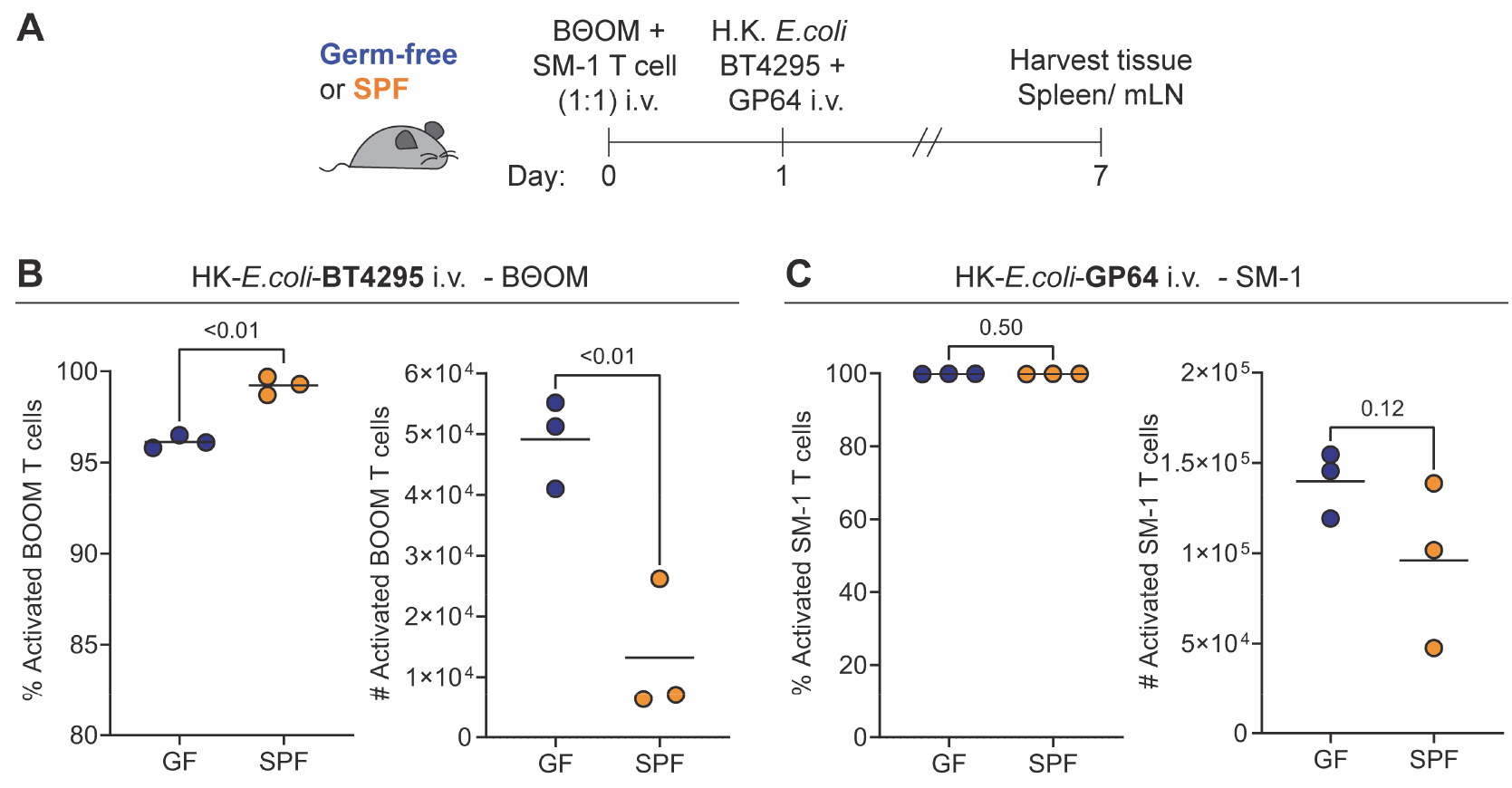
Systemic challenge with *E.coli*-BT4295 and *E.coli-GP64.* **(A)** Experimental set-up of *E.coli-*BT4295 and *E.coli-GP64* systemic challenge in GF and SPF mice. BΘOM and SM-1 T cells were transferred at a 1:1 ratio one day before challenge with inactivated *E.coli-*BT4295 and *E.coli-GP64.* **(B)** Percentage and number of activated BΘOM T cells in GF and SPF mice both in spleen. **(C)** Percentage and number of activated SM-1 T cells in GF and SPF mice both in spleen.

**Supplementary Figure 8:**
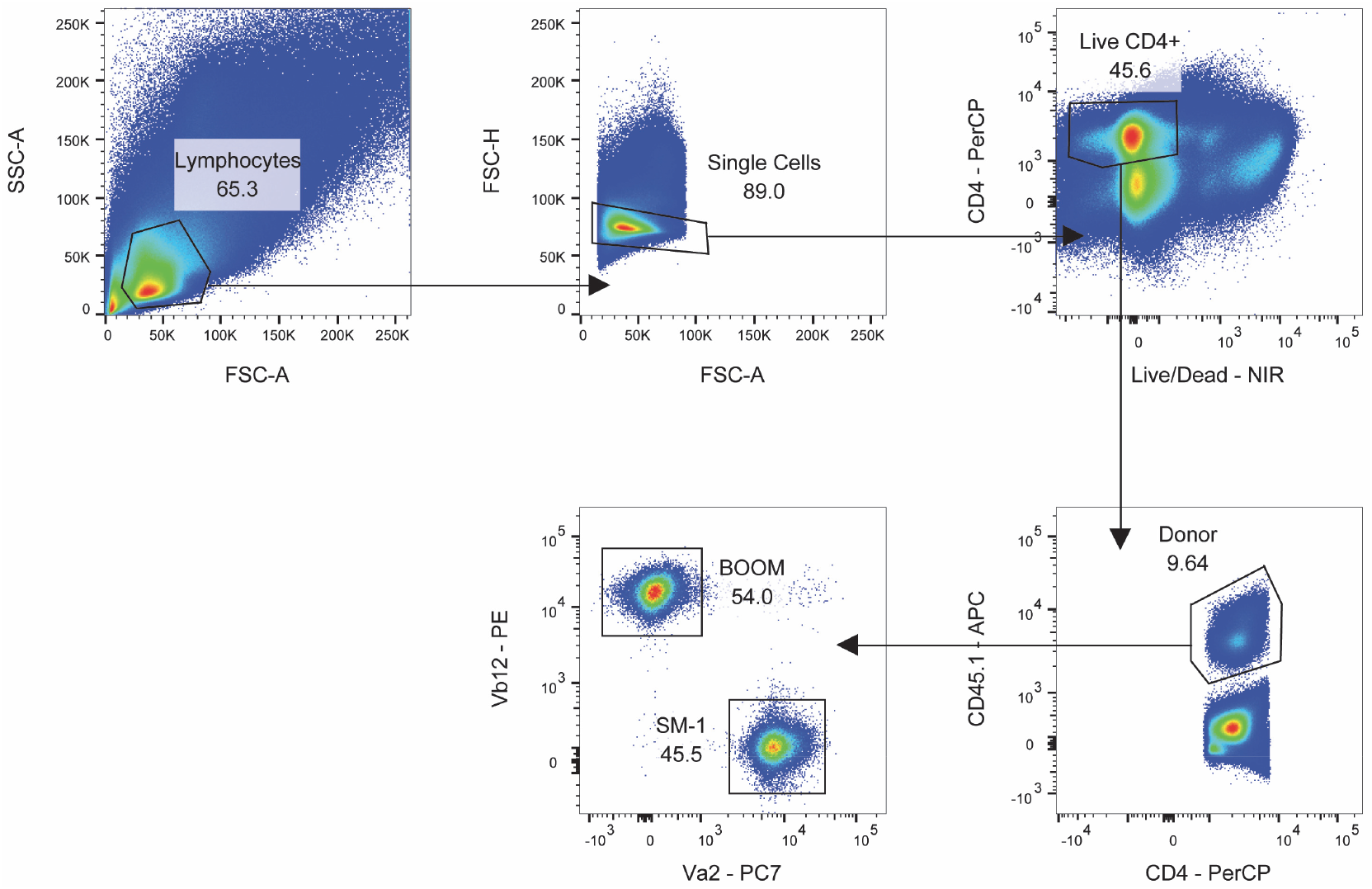
Gating strategy. After gating for live single-cell lymphocytes, donor CD4+ T cells were identified by the expression CD45.1. BΘOM and SM-1 T cells were gated by their expression of their corresponding TCR chain (Vβ12 for BΘOM and Vα2 for SM-1). Activated T cells were identified by CD44 and dilution of the CTV proliferation dye as in Supplementary Figure 1.

**Supplementary Figure 9:**
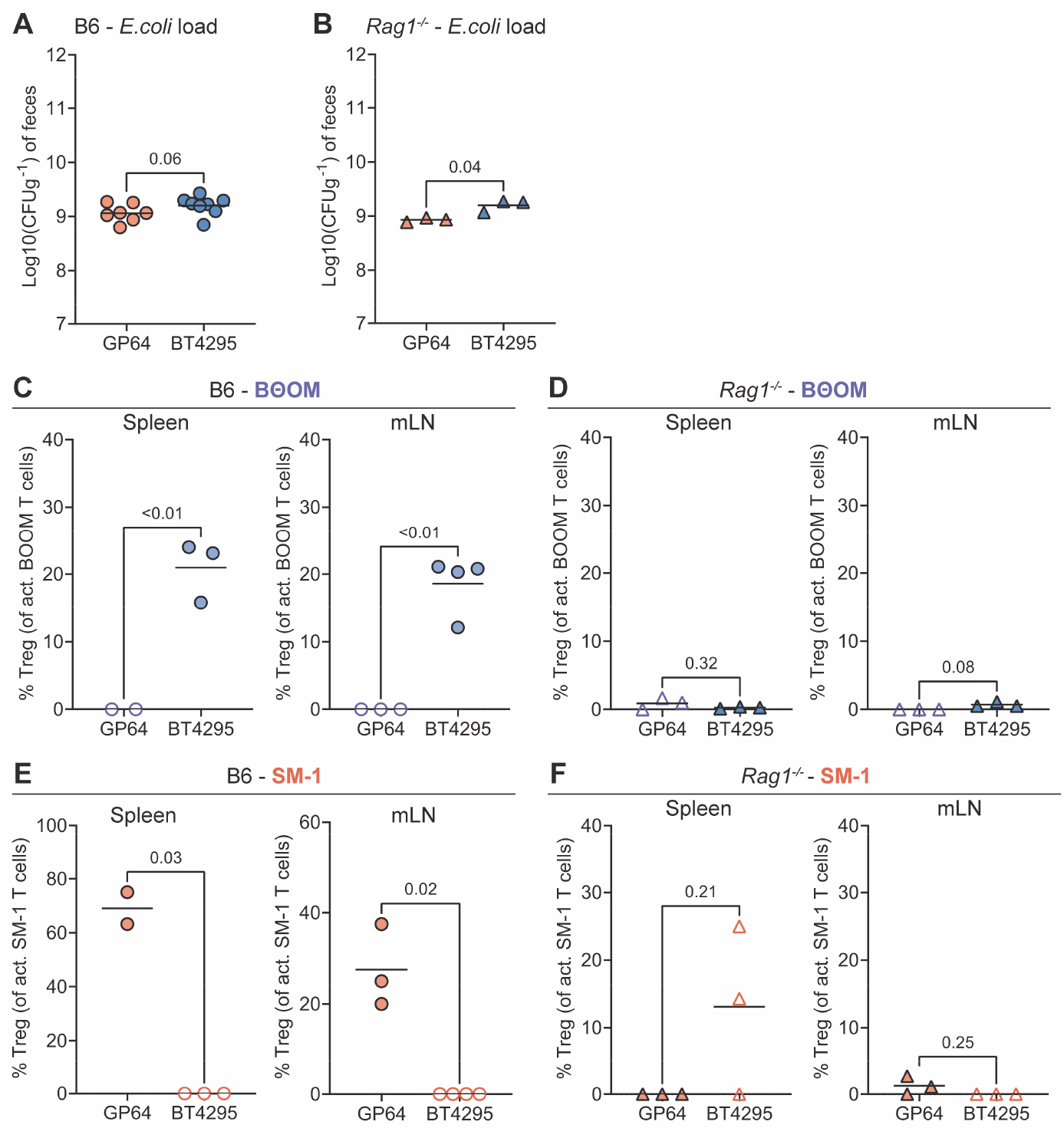
Bacterial load in gut lumen and induced regulatory T cells (Tregs) in BΘOM and SM-1 T cells. **(A and B)** *E.coli* strain luminal load in feces at the day of adoptive T cell transfer in **(A)** B6 and **(B)** *Rag1^-/-^.* **(C-D)** Percentage of Tregs among activated BΘOM T cells in **(C)** B6 and **(D)** *Rag1^-/-^.* **(E-F)** Percentage of Tregs among activated SM-1 T cells in **(C)** B6 and **(D)** *Rag1^-/-^.*

**Supplementary Figure 10:**
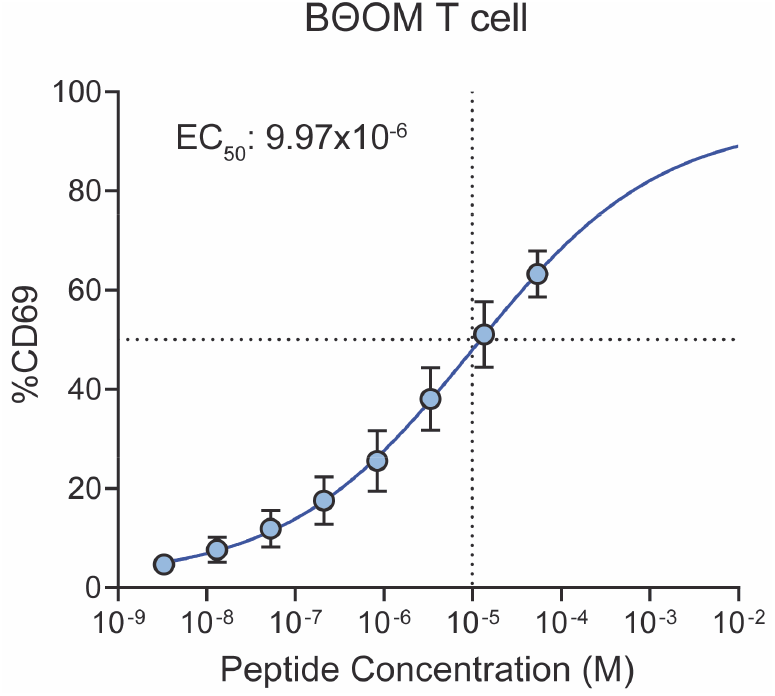
Peptide dose-activation curves. BΘOM T cells were cultured overnight with BT4295 peptide-pulsed splenocytes. The percentage of CD69+ BΘOM T cells is used as a readout for activation. (n=30 per dilution, each dot represents mean and bars standard deviation)

